# Balancing dynamic tradeoffs drives cellular reprogramming

**DOI:** 10.1101/393934

**Authors:** Kimberley N. Babos, Kate E. Galloway, Kassandra Kisler, Madison Zitting, Yichen Li, Brooke Quintino, Robert H. Chow, Berislav V. Zlokovic, Justin K. Ichida

## Abstract

Although cellular reprogramming continues to generate new cell types, reprogramming remains a rare cellular event. The molecular mechanisms that limit reprogramming, particularly to somatic lineages, remain unclear. By examining fibroblast-to-motor neuron conversion, we identify a previously unappreciated dynamic between transcription and replication that determines reprogramming competency. Transcription factor overexpression forces most cells into states that are refractory to reprogramming and are characterized by either hypertranscription with little cell division, or hyperproliferation with low transcription. We identify genetic and chemical factors that dramatically increase the number of cells capable of both hypertranscription and hyperproliferation. Hypertranscribing, hyperproliferating cells reprogram at 100-fold higher, near-deterministic rates. We demonstrate that elevated topoisomerase expression endows cells with privileged reprogramming capacity, suggesting that biophysical constraints limit cellular reprogramming to rare events.

## Introduction

Cellular reprogramming through forced overexpression of transcription factor cocktails redirects the transcriptional state of a cell, converting its fate into an increasing list of lineages. By providing access to rare, inaccessible cell types in unique genomic contexts, reprogramming massively expands the potential for in vitro disease modeling. Investigations into neurological diseases, previously limited by the supply of relevant human cells and the weak fidelity of murine models, have advanced through cellular reprogramming. Insights into the molecular changes that occur in disease contexts are already generating novel therapeutic strategies (1-3). In addition, because direct lineage conversion can preserve epigenetic signatures derived from the starting cells, this approach has the potential to lead to a better understanding of enigmatic processes such as aging (4-6).

However, cellular reprogramming remains a rare event (7). Although lineage conversion into somatic lineages is becoming increasingly desirable for translational studies, efforts to identify mechanisms that limit transcription factor-mediated lineage conversion have focused primarily on iPSC generation (8-15). Identified mechanisms appear to be specific to iPSC reprogramming, such as the identification of a rapidly-cycling pre-iPSC intermediate (10,12) or direct interactions of epigenetic complexes with Oct4, Sox2, and Klf4 (16,17). Indeed, inclusion of epigenetic modifiers such as valproic acid and trichostatin A increase reprogramming to iPSCs, but has limited utility in other lineage conversions (18-20). Even rapidly cycling and other “privileged” cells form iPSCs non-deterministically, indicating that there are critical reprogramming determinants that remain unidentified (8,10). For these reasons, it is unclear what mechanisms limit direct lineage conversion more broadly.

Roadblocks limiting direct lineage conversion have practical implications. In particular, post-mitotic cell types require a high efficiency of conversion in order to have translational utility. Generating mature and homogeneous cultures of a target cell type remains a key barrier limiting disease modeling and cell transplantation studies (21-24). Moreover, the particular methods of cellular reprogramming can leave distinct transcriptional and functional vestiges within the final cell types. For example, previous studies have shown that directly reprogrammed cells may retain expression of gene regulatory networks associated with the fibroblasts from which they originate (25,26).

We sought to identify universal roadblocks to reprogramming that extend beyond iPSCs into other lineages and define strategies to overcome them. To this end, we examined systems-level constraints limiting the conversion of fibroblasts into motor neurons as well as other paradigms. We find that during lineage conversion, addition of the reprogramming factors induces a sharp increase in the rate of transcription in cells and significantly reduces the percentage of fast-cycling cells, highlighting the existence of tradeoffs between transcription and cell replication during the conversion process. Most cells display either a high rate of transcription and limited proliferation or a high rate of proliferation and limited transcription, with both cell states being largely refractory to reprogramming, However, we identify a privileged population of cells capable of both fast-cycling and high transcription rates that contribute to the majority of reprogramming events. Using genetic and chemical factors, we expand the hypertranscribing, hyperproliferating cell (HHC) population and achieve induced motor neuron reprogramming at near-deterministic rates. Through transcriptional profiling, we identify topoisomerases, enzymes that curate DNA supercoiling introduced during transcription and DNA replication, as key regulators that support the emergence and expansion of these privileged HHCs. We also show that by expanding the population of HHCs, we accelerate the maturation and reduce the heterogeneity of the resulting cells. Our results suggest that dynamic biophysical constraints limit cellular reprogramming to rare events.

## Hyperproliferation promotes conversion to multiple, post-mitotic fates

Previous work identifying cellular features that enable cellular reprogramming examined the role of proliferation and found that fast-cycling, or hyperproliferating cells preferentially reprogrammed into induced pluripotent stem cells (iPSCs) (10). However, because iPSCs are necessarily fast-cycling, these studies could not determine if hyperproliferation is generally required for transcription factor-mediated reprogramming or if it is specifically required in iPSC generation because it aligns the cell division rate of the starting somatic cells with that of pluripotent stem cells (10,11). To more generally examine the transient role of hyperproliferation during conversion, we investigated the impact of hyperproliferation on fibroblasts driven to a post-mitotic fate. First, we defined “hyperproliferation” (a.k.a. “fast-cycling”) as a two-fold increase in division rate above population average, which is consistent with published studies (10). We focused on the motor neuron lineage because it is a well-defined neuronal subtype with established markers and reporters. Utilizing mouse embryonic fibroblasts (MEFs) isolated from Hb9::GFP transgenic mice, we generated induced motor neurons (iMNs) by viral overexpression of six transcription factors (Ascl1, Brn2, Myt1l, Ngn2, Isl1, Lhx3) (6F) as described previously (7).

To examine the effect of proliferation on conversion into iMNs, we applied genetic perturbations to increase the rate of cell division and quantified the rate of iMN reprogramming by counting Hb9::GFP positive cells within a fixed number of seeded cells. To track the effect of each perturbation on cell proliferation, we labeled cells with the stable dye CFSE 24 hours after infection with the transcription factors (Fig. 1A). Dilution of the CFSE dye occurs via cell division, resulting in dim, fast-cycling cells and brightly-stained, slowly dividing cells (Fig. 1A). We quantified CFSE levels by flow cytometry 72 hours post-label (4 days post-infection (dpi)) (Fig. 1A). Overexpression of the reprogramming factors reduced the percentage of fast-cycling cells ten-fold compared to uninfected fibroblasts (Fig. 1B). Transducing with DsRed retrovirus did not elicit this change, suggesting that this effect was dependent on the transgenic factors being transcription factors (Fig. S1A). Moreover, the effect was not specific to neuronal transcription factors, as overexpression of the iPSC reprogramming factors Oct4, Sox2, and Klf4 similarly reduced the percentage of fast-cycling cells (Fig. S1B, C). Further reducing the rate of cell division via overexpression of p21 or mitomycin C (MMC) treatment significantly lowered iMN conversion (Fig. S2A, B). In contrast, increasing the population of fast-cycling cells through inclusion of p53DD (DD), a p53 mutant lacking a DNA-binding domain (27), resulted in significantly higher conversion (Fig. 1C). We did not observe reduced rates of apoptosis in DD conditions, eliminating inhibition of apoptosis as the principle mechanism of increased conversion with DD (Fig. S2C).

**Figure 1.**
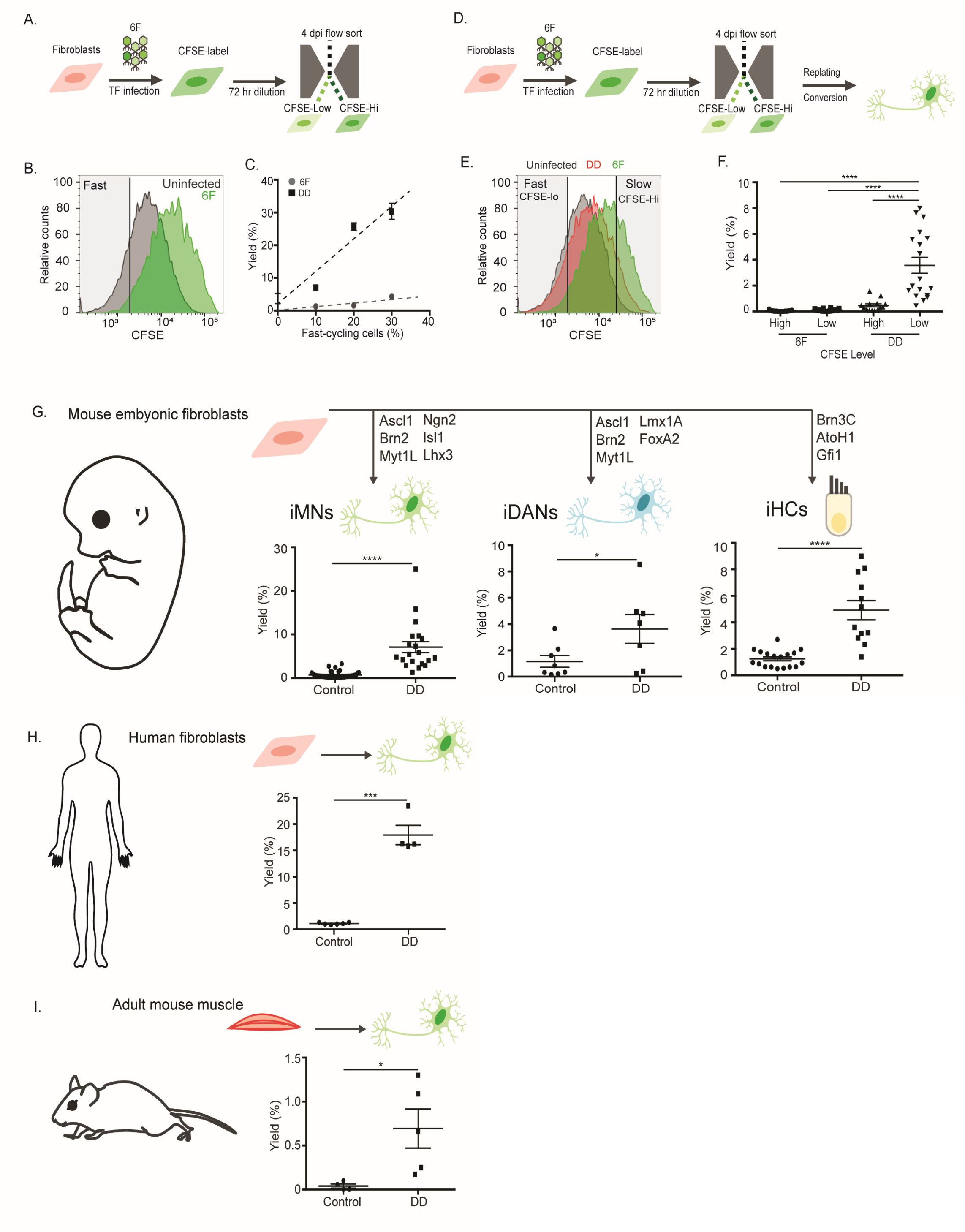
Hyperproliferation promotes conversion to a post-mitotic fate. (A) Schematic of CSFE-based sorting during reprogramming and reprogramming quantification assays. (B) Representative histograms of CFSE intensity for uninfected cells and 6 factor (6F) infected cells measured by flow cytometry at 4 dpi. (C) Conversion yield plotted over population of fast cycling cells (% of total viable cells) measured by flow cytometry. Population of fast-cycling cells was determined by gating cells with CFSE intensity 8-fold lower than mean CFSE intensity of uninfected, indicating 2-fold increase in division rate (e.g. expected to be approximately three additional cell divisions over 72 hrs based on 24 hr average cell cycle rate). Conversion yield determined by counting iMNs (Hb9::GFP+ cells with neuronal morphology) divided by the number of cells seeded for conversion. (D) Schematic of CSFE-based sorting and replating of populations for reprogramming quantification assays. (E) Representative histograms of CFSE intensity for uninfected cells, 6 factor (6F), and DD conditions by flow cytometry at 4 dpi with gates showing CFSE-Low (e.g. fast-cycling cells) and CFSE-Hi (e.g. slow cycling cells). (F) Yield of iMNs from 6F or DD reprogramming populations sorted by CFSE-intensity (e.g. CFSE-Low and CFSE-Hi) at 4 dpi. Percent yield determined by counting total iMNs generated normalized by total number of cells counted per population at 4 dpi. (G) Yield of iMNs, iDAs, and iHCs with defined transcription factor cocktails (Control) compared to yield in presence of p53DD (DD). (H) Yield of iMNs generated from human fibroblasts with factors alone (Control) compared to yield in presence of p53DD (DD). (I) Yield of iMNs generated from adult mouse muscle explants with factors alone (Control) compared to yield in presence of p53DD (DD). Data presented as mean + SEM of at least three biological replicates. *p ≤ 0.05, **p ≤ 0.01, * **p ≤ 0.001, and ****p ≤ 0.0001.

To definitively determine if fast-cycling cells convert into iMNs at higher rates than slow-cycling cells, we prospectively isolated fast- and slow-cycling cells from converting cultures by flow cytometry 72 hours after CFSE labeling (at 4 dpi), replated, and measured their ability to form iMNs at 14-17 dpi (Fig. 1D-F). We identified 4 dpi as important inflection point in the reprogramming process because induction of DD expression after 4 dpi shows limited efficacy to increase reprogramming, indicating DD mediates critical processes in the first four days of reprogramming (Fig. S2D). Prospectively-isolated fast-cycling cells converted to iMNs more efficiently than slow-cycling cells, supporting the notion that hyperproliferation promotes reprogramming into non-dividing cell types as well as iPSCs. However, this effect was very small in the absence of DD (Fig. 1F). These results indicate that while high proliferation rates can promote conversion into iMNs, reprogramming requires an alternative positive determinant of cellular reprogramming provided by DD activity.

To examine the generality of the hyperproliferation-mediated reprogramming boost in the DD condition, we employed different protocols to generate an array of post-mitotic cell types. Inclusion of DD increased reprogramming of MEFs into induced dopaminergic neurons (iDANs) via Ascl1, Brn2, Mytl1L, LmxA1, and FoxA2 (28), and induced inner ear hair cells (iHCs) via Atoh1, Gata3, and Brn3C (Fig. 1G) (29). Additionally, the reprogramming increase extended across species and age in the starting cell populations to include conversion of human adult and neonatal fibroblasts into iMNs (Fig. 1H and Fig. S3A), and mouse adult myoblasts (30) and tail tip fibroblasts into iMNs (Fig. 1I and Fig. S3B). These results indicate that DD increases the reprogramming potential of fast-cycling cells to promote the reprogramming of somatic cells from different ages and species into post-mitotic lineages.

## Hypertranscribing, hyperproliferating cells drive reprogramming

The extensive increase in reprogramming across protocols and species suggests a general mechanism for cell division promoting the full transcriptional realignment required for reprogramming. However, fast-cycling cells required DD to achieve significantly increased reprogramming rates. Consistent with previous studies on iPSC conversion (10), fast-cycling cells did not reprogram deterministically in our iMN conversions (Fig. 1C, F). This suggests that other cellular features play an important role in reprogramming. We found that factors previously shown to be capable of enabling deterministic iPSC reprogramming, such as Mbd3 depletion (9), did not significantly increase iMN conversion efficiency, suggesting that these factors could not account for the non-deterministic reprogramming efficiency observed in induced neuron conversion (Fig. S4).

Given that introduction of transcription factors, which we anticipated would increase transcription rates, sharply reduced replication (Fig. 1B), we wondered if cells frequently fail to sustain high levels of both transcription and replication in reprogramming, and whether this might cause conversion to stall. Utilizing 5-ethynyl uridine (EU) incorporation to measure transcription rate, we profiled cells over early time points for both cycling rate and transcription rate (Fig. 2AC). As we had anticipated, addition of the reprogramming factors increased the rate of transcription by 50% from 1 to 2 dpi (Fig. 2D), coinciding with the expression of factors at 2 dpi. Both in the presence or absence of transcription factor overexpression, we observed an inverse relationship between cycling and transcription rates (e.g. fast-cycling cells transcribed RNA at lower rates than slow-cycling cells)(Fig. 2C, E). However, the increased rate of transcription induced by transcription factor overexpression reduced the average rate of proliferation beyond that of uninfected cells (Fig. 2C). Therefore, there is an inverse relationship between cell replication and increased transcription rates induced by transcription factor overexpression.

**Figure 2.**
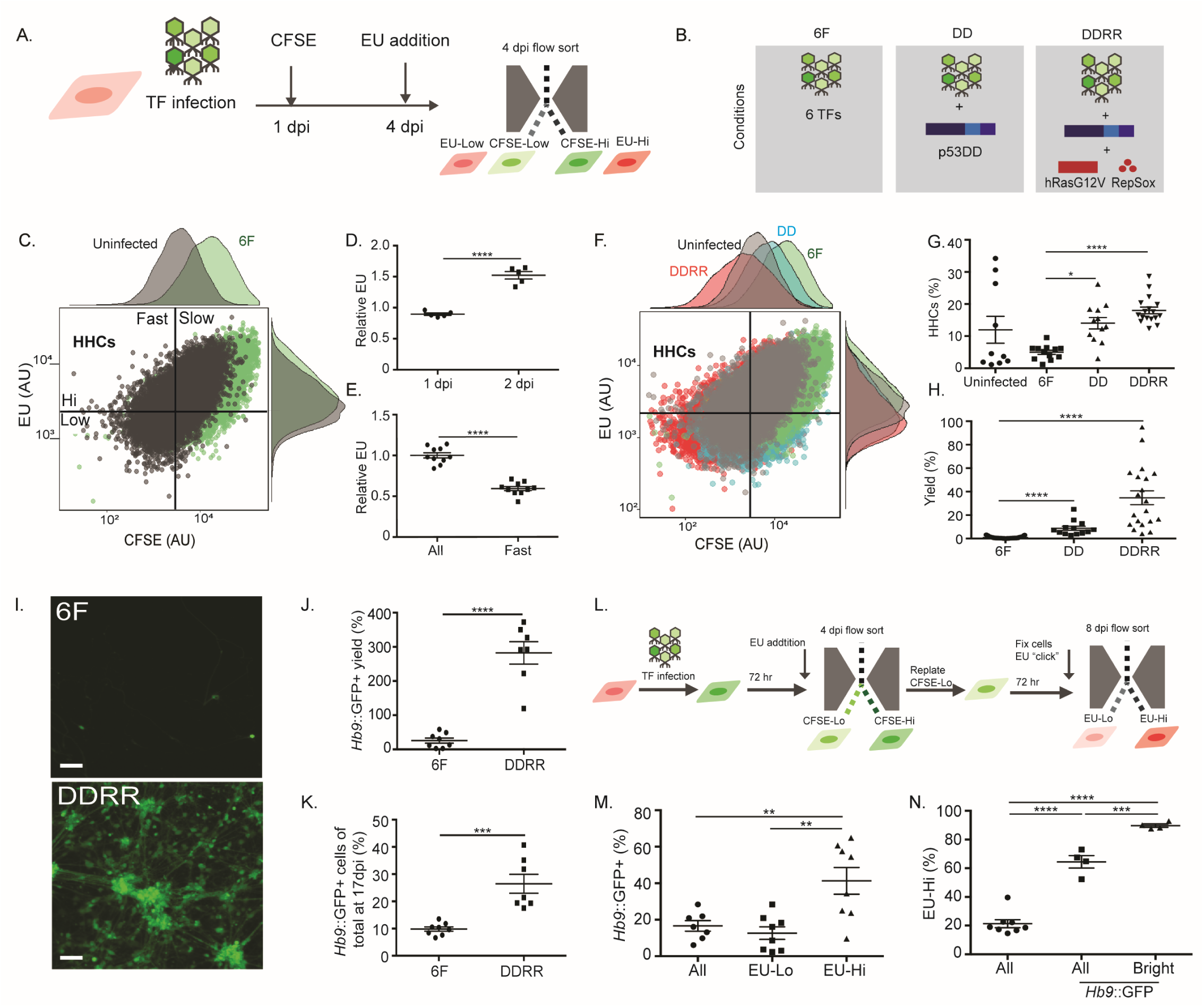
Hypertranscribing, hyperproliferating cells drive reprogramming. (A) Schematic of CFSE-EU assay for measuring transcription and proliferation rates in converting cells via flow cytometry at 4 dpi. Transcription rates measured through 5-ethynyl uridine (EU) incorporation during 1 hr incubation with 1mM EU, followed by “click” reaction with fluorescent dye to visualize EU incorporation. CFSE assay performed as described previously. (B) Legend of genetic and chemical combinations used for three primary conditions employed in conversion. 6F, 6 transcription factors only. DD, 6 transcription factors + p53DD. DDRR, 6 transcription factors + a combination of p53DD, hRasV12, a mutant of hRas, and RepSox, a TGF-β inhibitor. (C) Representative dot plot of CFSE intensity, and fluorescently labeled-EU intensity of uninfected MEFs (grey) and 6F infected cells (green) at 4 dpi via flow cytometry. Histograms of CFSE and EU intensity adjacent to dot plot. Quadrant to demark hypertranscribing, hyperproliferating cells (HHCs) set by reference to 6F condition. Fast and slow cycling cells set by selecting CFSE value 6F condition to allow the dimmest 15%. High EU values set by top half of 6F, resulting in ∼7% HHCs in 6F. AU defined as arbitrary units. (D) EU incorporation rate at 1 and 2 dpi in 6F-infected MEFs relative to non-infected control measured via flow cytometry. (E) EU incorporation rate of whole population (All) compared to fast cycling cells (Fast) measured in uninfected MEFs via flow cytometry at 4 dpi. (F) Representative dot plot of CFSE intensity and fluorescently labeled-EU for uninfected cells (UIC, grey), 6F (green), DD (blue), and DDRR (red). Histograms of CFSE and EU intensity adjacent to dot plot. (G) Fraction of the total population of HHCs for uninfected UIC, 6F, DD, and DDRR condition as assayed via flow cytometry at 4 dpi. (H) Yield of iMNs in 6F, DD, or DDRR conditions. (I) Representative images of 6F and DDRR-iMNs at 17 dpi taken at 10x magnification. Scale bars represent 100μm. (J) Percent yield of Hb9::GFP+ cells for 6F and DDRR conditions counted via flow cytometry at 17 dpi and normalized to number of seeded cells. (K) Fraction of Hb9::GFP+ cells for 6F and DDRR conditions normalized to total population of cells measured via flow cytometry at 17 dpi. (L) Schematic of CFSE-EU-pulse label assay to sort and label HHCs at 4 dpi followed by evaluation of Hb9::GFP intensity at 8 dpi. Cells were treated with CFSE at 1 dpi as previously described. At 4 dpi, cells were incubated with EU for 3 hrs prior to harvesting. Fast-cycling cells were gated and collected via FACS of CFSE intensity (e.g. CFSE-Low). Collected cells were replated and allowed 96 hours to convert. At 8 dpi, cells were harvested, fixed, and “clicked” to fluorescently label EU incorporated at 4 dpi. Cells were analyzed via flow cytometry to evaluate EU and Hb9::GFP intensity. (M) Fraction of Hb9::GFP+ cells in DDDR conditions for various gated populations. Cells gated for low EU intensity (EU-Lo, e.g. lowest three quartiles) and high EU-intensity (EU-Hi, e.g. highest quartile) at 8 dpi compared to all cells (All). (N) Fraction of replated fast-cycling cells in DDRR conditions gated for high EU intensity at 8 dpi. The whole population (All) contained 25% EU-Hi cells while Hb9+ and Bright cells (e.g. top half of all Hb9+ cells) had significantly larger fraction of EU-Hi cells. p ≤ 0.05, **p ≤ 0.01, * **p ≤ 0.001, and ****p ≤ 0.0001.

To determine if DD might enable fast-cycling cells to reprogram efficiently by providing them the ability to maintain a high rate of transcription, we measured cellular proliferation and transcription rates during reprogramming in the presence or absence of DD. In addition to restoring the population of fast-cycling cells, DD also increased the transcription rate of fast-cycling cells, resulting in a larger population of hypertranscribing, hyperproliferating cells (HHCs) and greater iMN yield (Fig. 2F-H). To determine if further increasing transcription in fast-cycling cells could improve reprogramming, we supplemented DD in two ways. First, we overexpressed a Ras mutant (hRasG12V) that was previously shown to globally increase transcription in human mammary epithelial cells (31). Second, we included treatment with RepSox, a small molecule inhibitor of TGF-β signaling, a pathway that has been shown to suppress transcription levels (32). This condition, termed DDRR (DD, Ras, RepSox) (Fig. 2B), significantly increased the population of HHCs in the presence of the six reprogramming factors (Fig. 2G) and resulted in a 100-fold increase in iMN yield (Fig. 2H, I). In DD conditions, RepSox and Ras cooperatively increased conversion (Fig. S5). To determine if the increased transcription rates in fast-cycling cells may explain the increased conversion efficiency in the presence of DDRR, we measured iMN reprogramming with or without the RNA polymerase inhibitor α-amanitin. Consistent with the notion that a high rate of transcription was required for the DDRR effect, reducing transcription by 20% at 4 dpi with α-amanitin treatment significantly reduced reprogramming (Fig. S6).

Given the density of Hb9::GFP+ cells in DDRR conditions, plate-based estimates of cell numbers represent an underestimate of yield. To more exhaustively quantify cell number, we employed flow cytometry. Quantifying the cell population for Hb9::GFP+ cells at the end of conversion, we analyzed total yield of Hb9::GFP+ cells based on starting number of cells and the Hb9::GFP+ fraction of the whole culture at 17 dpi. While the 6F alone condition resulted in fewer than 1 iMN per 100 MEFs plated, DDRR yielded about 200 iMNs per 100 MEFs plated (Fig. 2I, J), a 200-fold increase in yield. To take into account the fact that MEFs expand beyond the initial amount plated and determine the true efficiency of iMN conversion, we also quantified the percent of Hb9::GFP+ cells out of the total number of cells at the end of the conversion process. In the absence of DDRR, 90% of cells failed to activate Hb9::GFP. With DDRR, 30% of the population activated Hb9::GFP (Fig. 2K). Taking into account that HHCs represent approximately 20-30% of the whole population in DDRR conditions (Fig. 2G) and comprise the majority of reprogrammable cells (see Fig. 2L-N below), 30% of MEFs activating Hb9::GFP represents near-deterministic reprogramming from this population. These data suggest that we have increased the reprogrammable population by expanding the number of cells capable of exhibiting both hypertranscription and hyperproliferation.

While increasing the population of HHCs at early time points increased conversion rate, we sought to definitively determine if this particular fast-cycling population of cells possessed privileged reprogramming relative to slower transcribing, fast-cycling cells. To test if HHCs identified at 4 dpi possess greater reprogramming potential relative to hyperproliferating cells with lower transcription rates, we performed a reprogramming experiment with prospective labeling of HHCs and non-HHCs. We first flow-purified all fast-cycling cells that had been pulse-labeled with EU at 4 dpi (Fig. 2L). At 8 dpi, we fixed and “clicked” EU-labeled cells to measure EU incorporation. We identified HHCs by high EU levels (e.g. top quartile of EU intensity evaluated in all cells) and analyzed for Hb9::GFP expression.

Of the HHC population, over 40% expressed Hb9::GFP at 8 dpi, while only 13% of the non-HHC population expressed Hb9::GFP (Fig. 2M). Thus, HHCs were 3 to 5 times as likely to activate Hb9::GFP relative to hyperproliferative non-HHCs. Because our combined imaging/single cell qRT-PCR analyses had shown that the GFP intensity of Hb9::GFP+ cells was strongly correlated with neuronal morphology and gene expression (Fig. S7), we examined the EU intensity of bright Hb9::GFP+ cells (bright was defined as the top 50% of Hb9::GFP+). We observed that 90% of bright Hb9::GFP+ cells had high EU intensity (e.g. top quartile of EU intensity evaluated in all cells), meaning that of the Hb9::GFP+ cells that advanced to the terminal neuronal stage of reprogramming, the vast majority of them originated from HHCs (Fig. 2N). Thus, HHCs possess significantly greater reprogramming potential than non-HHCs, including cells that are hyperproliferative but do not hypertranscribe. Taken together, our data indicate that the inability of most cells to sustain hypertranscription and hyperproliferation limits reprogramming to rare cell populations. By expanding the population of cells capable of mediating both processes, we expand reprogramming to near-deterministic rates.

## Combined hypertranscription and hyperproliferation promotes both early and intermediate stages of iMN reprogramming

Previous analysis of gene regulatory networks (GRNs) identified that components of the fibroblast GRN remain active within induced neurons (iNs) (25,26). One interpretation of this finding is that induced neurons fail to adopt a fully neuronal transcriptional program. Alternatively, mechanisms that limit induced neuron reprogramming may arrest cells at intermediate states leading to heterogeneous cultures comprised of fully neuronal and partially neuronal cells.

Using a combination of live imaging and cross-sectional analysis across multiple endpoints, we identified a post-mitotic intermediate state characterized by Hb9::GFP reporter activation and retention of a fibroblast morphology (e.g. Hb9::GFP+ fibroblast (Fig. 3A, top panel)). This state frequently preceded Hb9::GFP+ iMN formation (Fig. S8A), and in iMN conversions using 6F alone, at least half of Hb9::G-FP+ cells remained trapped in this state (Fig. S8B).

**Figure 3.**
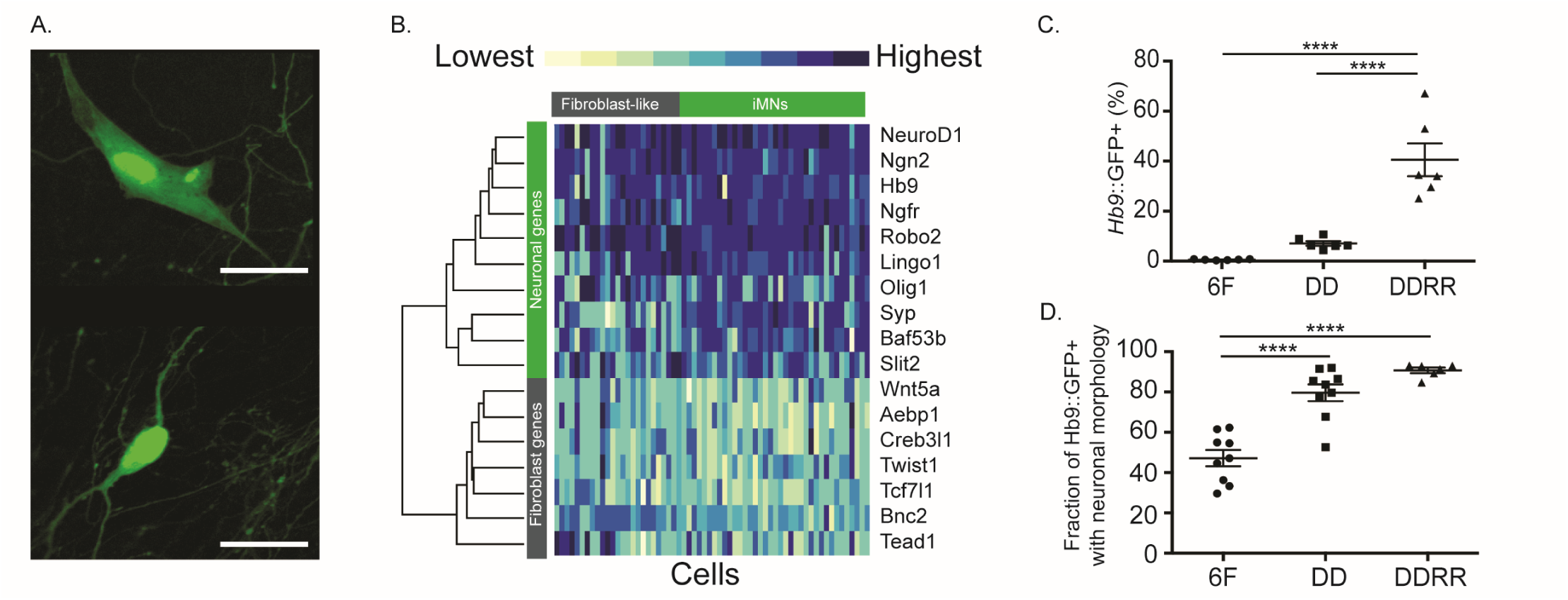
Cells with the capacity for both hyperproliferation and hypertranscription transition through the Hb9+ intermediate stage more efficiently. (A) Representative images of Hb9::GFP+ cells with fibroblast (top) or neuronal (bottom) morphology. Scale bars represent 20μm. (B) Heatmap of relative expression for single cells with either fibroblast (top gray, n=24) or neuronal (top green, n=39) morphology for qPCR assays for fibroblast (side gray) or neuronal (side green) genes. (C) Fraction of Hb9::GFP+ cells for 6F, DD, or DDRR conditions measured by flow cytometry at 8 dpi. (D) Fraction of Hb9::GFP+ with neuronal morphology of total Hb9::GFP+ cells for 6F, DD, or DDRR conditions at 17 dpi. Data presented as mean + SEM of at least three biological replicates. Significance determined by one-way ANOVA. ****p ≤ 0.0001.

To identify the transcriptional state associated with the Hb9::GFP+ intermediates, we isolated individual Hb9::GFP+ iMNs with a neuronal morphology and Hb9::GFP+ intermediates with fibroblast morphologies and measured gene expression via qRT-PCR. iMNs (Fig. 3A, bottom, 3B, top green) displayed increased expression of neuronal markers and decreased expression of fibroblast gene regulatory network (GRN) genes relative to Hb9::GFP+ fibroblast-like intermediates (Fig. 3B). Together, our data indicate that during induced motor neuron reprogramming, there is a molecular barrier that causes cells to accumulate in a partially-reprogrammed intermediate state possessing both neuronal and fibroblast gene expression. Therefore, previous studies may have detected fibroblast GRN transcription due to the presence of partially-reprogrammed intermediates rather than the induced neurons themselves retaining fibroblast properties (25,26). In addition, our analyses indicate that cellular morphology represents an important proxy for evaluating an individual cell’s transcriptional state during induced motor neuron conversion.

We hypothesized that by increasing the transcription rate of fast-cycling cells (i.e. the HHC phenotype), DD and DDRR might accelerate the transition from Hb9::GFP+ intermediates to iMNs with full adoption of the motor neuron transcriptional state. To test this, we evaluated the two rate-limiting steps of reprogramming, the activation of Hb9::GFP and morphological remodeling to iMNs, via longitudinal tracking across the two-week window of conversion. In the presence of DD, Hb9::GFP+ intermediates were four times more likely to adopt a neuronal morphology and fully convert into iMNs (Fig. S8C). To confirm these results, we counted Hb9::GFP+ cells by flow cytometry to determine the rate of Hb9::GFP activation at 8 dpi and the morphology of Hb9::GFP+ cells at 17 dpi for the 6F, DD, and DDRR conditions (Fig. 3C, D). Similar to our longitudinal tracking data, the rate of Hb9::GFP activation and iMN generation correlates with population size of HHCs. With 6F alone, less than 1% of cells activate Hb9::GFP, while 8% and 40% activate Hb9::GFP in the DD and DDRR conditions, respectively (Fig. 3C). Additionally, 50% of Hb9::GFP cells remained trapped in an intermediate state with 6F alone (Fig. 3D). In contrast, addition of DDRR to the 6F cocktail resulted in 90% of Hb9::GFP+ cells becoming iMNs at 17 dpi (Fig. 3D). These data indicate that DDRR does not increase the number of iMNs simply by expanding MEFs prior to reprogramming or preventing cell death during reprogramming. Instead, DDRR enables cells to maintain hypertranscription and hyperproliferation, which leads to efficient activation of Hb9::GFP and the systematic transition of partially-reprogrammed intermediates to the neuronal state.

## Topoisomerase expression enables simultaneous hypertranscription, hyperproliferation in HHCs

To identify the mechanisms that enable combined hypertranscription and hyperproliferation, we transcriptionally profiled single cells at different time points in conversion (Fig. 4A). To focus our analysis on cells that were on a successful reprogramming trajectory, we collected fast-cycling cells at 4 dpi (CFSE-Lo) and Hb9::GFP+ cells at 8 dpi (Fig. 4A).

**Figure 4.**
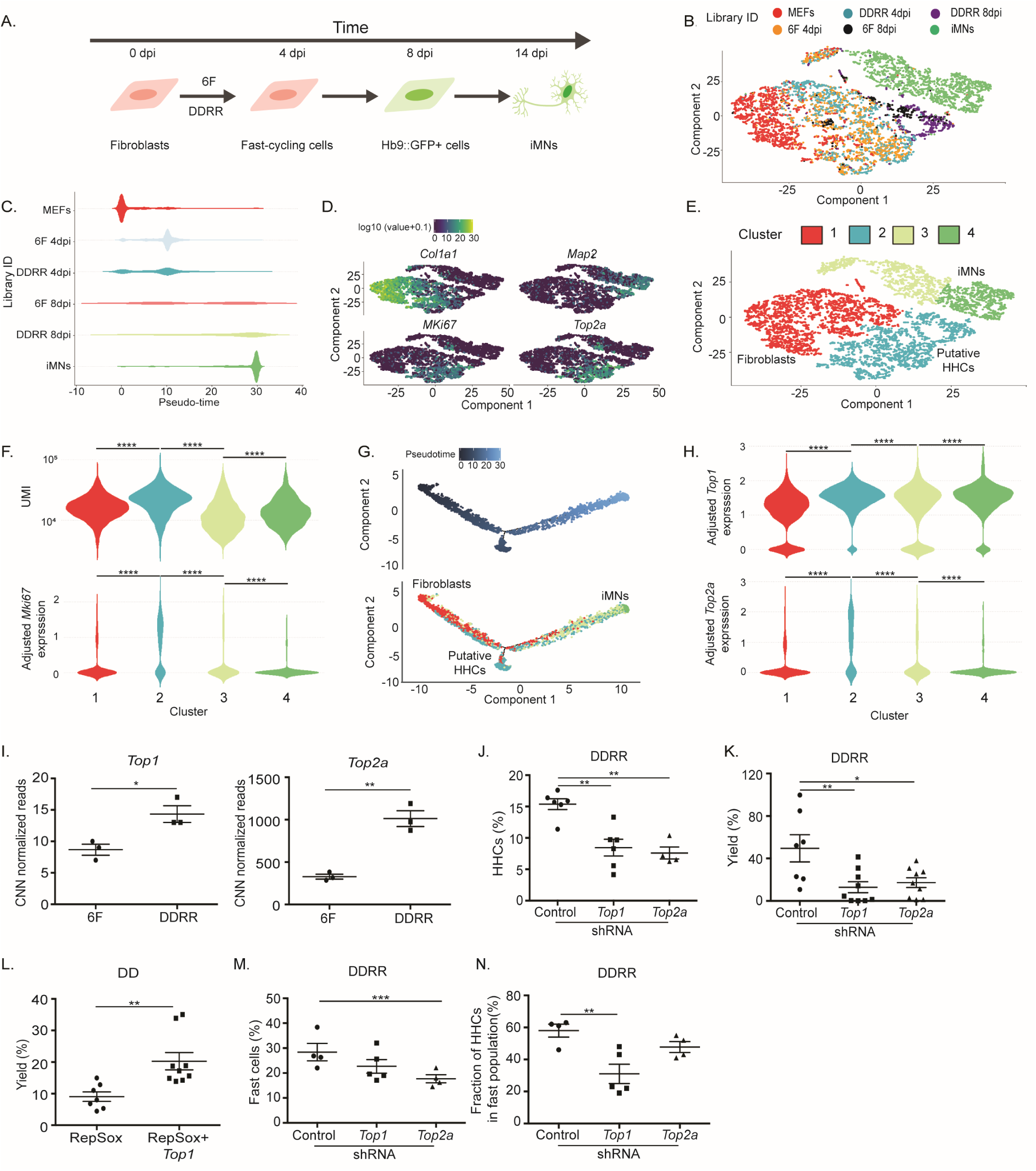
Topoisomerase expression enables cells to exhibit both hyperproliferation and hypertranscription. (A) Schematic of populations collected across conversion and profiled via single-cell RNAseq. Individual libraries were prepared for MEFs (1357), fast-cycling cells (e.g. CFSE-Low) for 6F (1174) and DDRR (1189) collected at 4 dpi (e.g. 6F 4dpi, DDRR 4dpi), Hb9::GFP+ cells for 6F (259) and DDRR (406) at 8 dpi (e.g. 6F 8dpi, DDRR 8dpi) and DDRR iMNs (1598) at 17 dpi (iMNs). (B) tSNE projection of all cells mapped during reprogramming colored by individual library. (C) Distribution of pseudotime across cells in each library population. (D) Relative expression colored by intensity of Cola1, Map2, Mki67, and Top2a over the populations in the tSNE. (E) Clustering of populations within the tSNE projection. (F) Violin plot of UMI (top, unique molecular identifiers) and relative Mki67 expression (bottom) for clusters identified in (D). (G) Reprogramming trajectory constructed through pseudotemporal ordering of single cells via Monocle orders cells from MEFs (red) to iMNs (green). (H) Violin plot of relative expression of Top1 (top) and Top2a (bottom) for clusters identified in (D). (I) Relative expression of Top1 (left) or Top2a (right) in 6F or DDRR conditions at 4 dpi in fast-cycling cells based on number normalization (CNN) RNAseq (J) Fraction of HHCs in population of converting cells in DDRR conditions treated with Control, Top1, or Top2a shRNAs. (K) Yield of iMNs in DDRR conditions treated with Control, Top1, or Top2a shRNAs. (L) Yield of iMNs in DD conditions treated with RepSox with or without Top1 overexpression. (M) Fraction of fast cells in total population as measured by flow cytometry at 4 dpi of DDRR cells treated with Control, Top1, or Top2a shRNAa. Significance analyzed with two-sided Student’s t-test. (N) Fraction of HHCs in population of fast-cycling cells as measured by flow cytometry at 4 dpi of DDRR cells treated with Control, Top1, or Top2a shRNAa. Except where otherwise stated, data presented as mean + SEM of at least three biological replicates. Significance determined by one-way ANOVA for multiple comparisons or a two-sided Student’s t-test for comparison of control and test. Significance summary: p > 0.05 (ns), *p ≤ 0.05, **p ≤ 0.01, ***p ≤ 0.001, and ****p ≤ 0.0001

To determine if cells in 6F and DDRR conditions take similar or distinct trajectories from the fibroblast to the iMN state, we profiled cells from both conditions (Fig. 4A, B). Analysis of tSNE clustering indicated that converting cells at 4 dpi and 8 dpi mapped between fibroblasts and fully-converted iMNs (Fig. 4B). As expected, cells at 4 dpi clustered closer to fibroblasts and Hb9::GFP+ cells at 8 dpi clustered closer to iMNs (Fig. 4B). At 4 dpi, cells with 6 factors alone mapped to similar locations as cells in DDRR conditions (Fig. 4B). However, at 8 dpi, most DDRR cells mapped to a location closer to iMNs, while the 6F population was split between this location and a location similar to the 4 dpi cells (Fig. 4B). These results suggest that converting cells traverse a similar trajectory in 6F and DDRR conditions.

Pseudotime analysis indicated that at 8 dpi, cells in DDRR conditions were much closer to the iMN state than cells in the 6F alone condition (Fig. 4C, note that the color scheme is consistent amongst C, E-H, and this scheme is distinct from that used in B). While Hb9::GFP+ cells at 8 dpi in 6F conditions bifurcated into two populations (correlating with Hb9::GFP+ fibroblasts and Hb9::GFP+ neurons) as expected from single-cell tracking and qRT-PCR data (Fig. 3A, B and 4C), DDRR cells showed a unimodal distribution aggregating near the iMN state (Fig. 4C), indicating that DDRR cells activating Hb9::GFP systematically go on to complete reprogramming. Together, these results suggest that cells traverse through a conserved trajectory during lineage conversion regardless of condition, but DDRR increases the speed and efficiency of reprogramming.

We next examined the different single cell clusters to identify the transcriptional programs that enable combined hypertranscription and hyperproliferation. As expected, fibroblasts expressed the highest levels of collagen genes such as Col1a1, while converting cells decreased collagen gene expression during transit to iMNs (Fig. S9A). Map2, a marker of post-mitotic neurons, was expressed by iMNs and a fraction of Hb9::GFP+ cells, but not fibroblasts (Fig. 4D, E). Converting cells grouped within two clusters. Cluster 1 remained close to the starting fibroblasts with low expression of proliferative genes such as Mki67 (Fig. 4D-F and Fig. S9B) and low total mRNA levels as measured by unique molecular identifiers (UMIs) (Fig. 4F). Cluster 2 contained cells with a proliferative signature including high expression of Mki67 (Fig. 4D), and high total mRNA levels (Fig. 4F), signifying hypertranscription. The increase in total mRNA levels in Cluster 2 cells was highly significant in comparison to the other cell clusters, which showed a lower distribution of UMIs per cell across libraries (Fig. 4F and S9C, D). Pseudotemporal ordering of clustered cells placed the Cluster 2/HHCs further ahead in reprogramming compared to Cluster 1 cells and on a separate branch, suggesting that the transcriptome of Cluster2/HHCs is highly divergent from that of non-HHCs (Fig. 4G). Thus, the transcriptional profile of Cluster 2 cells is consistent with the properties of the HHC population, which sustain combined hypertranscription and hyperproliferation. In the Cluster 2/HHC population, we identified increased expression of two topoisomerases (Fig. 4H). Top1 expression increased through reprogramming, while Top2a showed transient expression, peaking as cells transitioned from a fibroblast (high Col1a1, low Map2) state to an iMNs (low Col1a1, high Map2) state (Fig. S9A). Cell number normalized RNA-seq analysis comparing hyperproliferating cells in 6F and DDRR conditions at 4 dpi indicated that DDRR significantly increased levels of Top1 and Top2a (Fig. 4I). Topoisomerases remove supercoils induced by transcription and replication as well as resolve collisions between transcription and replication machinery (33,34). Therefore, we hypothesized that increased topoisomerase expression may allow HHCs to sustain high rates of replication and transcription.

To examine the necessity of each topoisomerase in supporting the HHC population and reprogramming, we introduced shRNAs targeting either Top1 or Top2a. While reduction of either topoisomerase reduced cellular viability (Fig. S10A), topoisomerase reduction significantly reduced the population of HHCs in the remaining cells under DDRR conditions (Fig. 4J). Consistent with HHCs comprising the majority of the reprogramming-competent cell population, knockdown of either topoisomerase also resulted in a significant drop in iMN yield in the DDRR condition (Fig. 4K). To test the sufficiency of Top1 to increase reprogramming, we overexpressed Top1 during conversion. Consistent with Top1 playing a key role in reprogramming, the addition of Top1 significantly increased iMN conversion (Fig. 4L). These results suggest that the upregulation of topoisomerases by DDRR is a critical mechanism by which these conditions expand the HHC population and enable highly efficient reprogramming.

We noticed that the full impact of Top1 overexpression in iMN conversion required the presence of p53DD and RepSox (Fig. 4L), which induce Top2a expression and increase the hyperproliferative cell population during reprogramming (Fig. S10B). To determine why Top1 may require p53DD and RepSox to affect reprogramming, we examined changes in the HHC population at 4 dpi induced by shRNA knockdown of either Top1 or Top2a. Interestingly, topoisomerase knockdown affected the loss of HHCs by different mechanisms. Knockdown of Top2a reduced the percentage of fast-cycling cells (Fig. 4M) without changing the distribution of transcription rates of fast-cycling cells (Fig. 4N). Conversely, Top1 knockdown reduced the transcription rate of fast-cycling cells with only a minor effect on the percentage of fast-cycling cells in the population. Therefore, our data suggest that Top1 requires p53DD and RepSox to exert its effect on reprogramming because Top1 mainly induces hypertranscription, whereas Top2a, which is increased by p53DD and RepSox, drives hyperproliferation. Taken together these data suggest that topoisomerases enable high levels of simultaneous transcription and replication in HHCs through distinct mechanisms. Increasing the expression of both topoisomerases via DDRR balances replication and transcription to promote the HHC state and cellular conversion.

## Converting HHCs adopt the induced motor neuron transcriptional program, accelerating maturation

Given the robust increase in reprogramming upon HHC induction with DDRR, we sought to determine the generality of inducing this HHC population in other reprogramming schemes. Compared to control or DD conversion cocktails, DDRR significantly enhanced conversion to several post-mitotic cell types including induced dopaminergic neurons and inner ear hair cells (Fig. 5A-C). These data suggest that expanding the HHC population may similarly enhance the utility of other reprogramming paradigms for high-throughput disease modeling and mechanistic studies.

**Figure 5.**
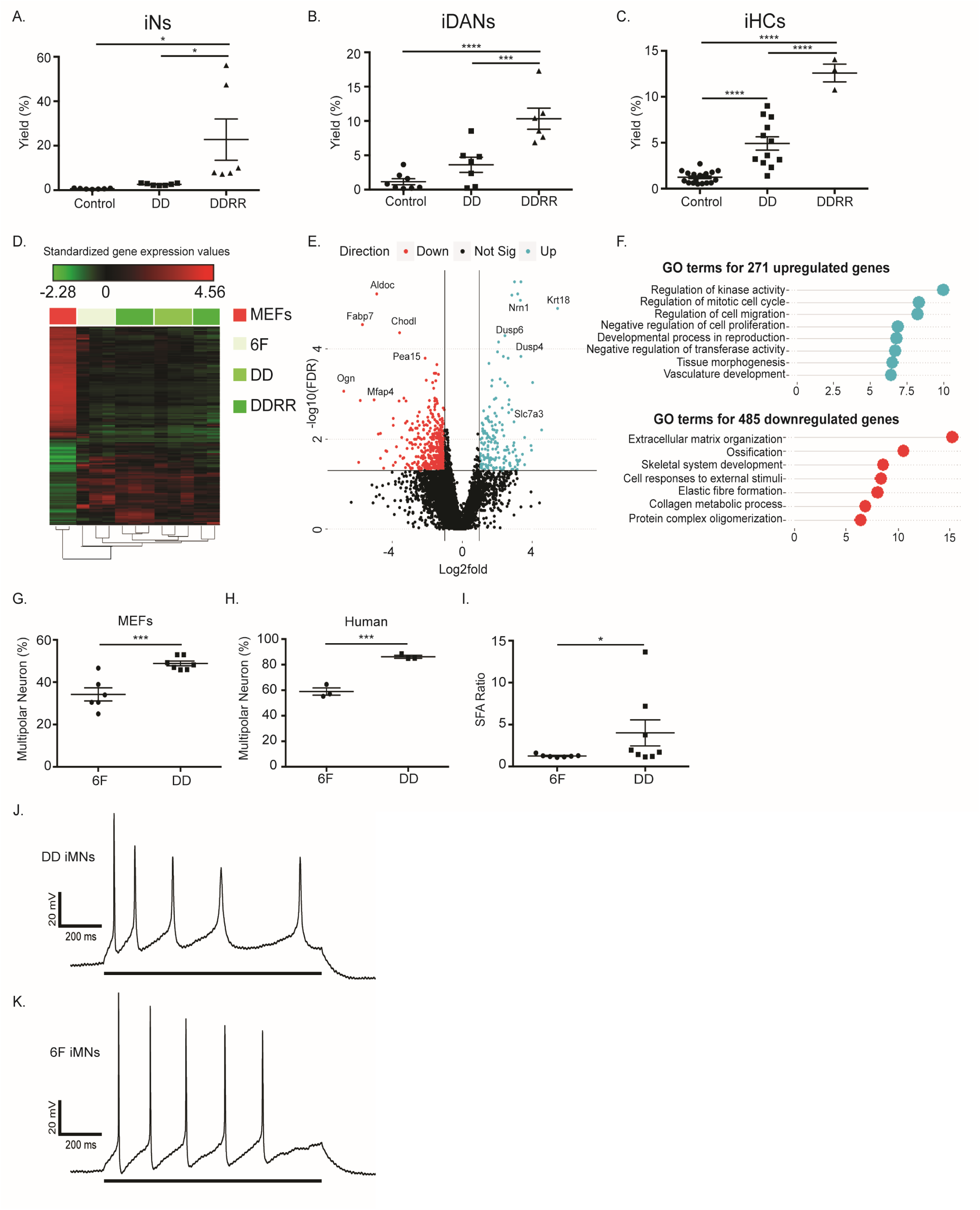
Converting HHCs adopt the induced motor neuron transcriptional program and accelerate morphological maturation. (A) Yield of induced neurons (iNs) for different conditions including control with factors only (e.g. Brn2, Ascl1, Myt1l), DD, and DDRR counted by MAP2+ cells at 17 dpi over number of cells seeded. (B) Yield of induced dopaminergic neurons (iDANs) for different conditions including control with factors only (e.g. Brn2, Ascl1, Myt1l, Lmx1A, FoxA2), DD, and DDRR counted by MAP2+ cells at 17 dpi. (C) Yield of induced inner ear hair cells (iHCs) for different conditions including control with factors only (e.g. Brn3C, Atoh1, Gfi1), DD, and DDRR counted by Atoh1::nGFP+ cells at 17 dpi. (D) Heat map of RNAseq collected from Hb9::GFP+ cells from different conditions on 17 dpi compared to starting MEFs across 1186 DEGs that vary between MEFs and Hb9::GFP+ cells. (E) Volcano plot comparison of genes up(blue) or downregulated (red) in DDRR versus 6F Hb9::GFP+ cells. (F) List of gene ontology (GO) terms for genes upregulated (top, blue) or downregulated (bottom, red) in DDRR cells compared to 6F. (G) Fraction of multipolar neurons for iMNs derived from MEFs in 6F and DD conditions. (H) Fraction of multipolar neurons for iMNs derived from primary human fibroblasts in 7F and DD conditions. (I) SFA ratio evoked APs of iMNs in 6F and DD conditions. Significance analyzed with Mann-Whitney log rank test. (J-K) Representative action potentials evoked in iMNs by a positive current injection (indicated by solid bar across bottom) illustrating SFA over the course of the stimulus of iMNs in DD (J) and 6F (K) conditions. Unless otherwise stated, data presented as mean + SEM of at least three biological replicates. Significance determined by one-way ANOVA for multiple comparisons or a two-sided Student’s t-test for comparison of control and test. Significance summary: p > 0.05 (ns), *p ≤ 0.05, **p ≤ 0.01, ***p ≤ 0.001, and ****p ≤ 0.0001.

To determine if the genetic and chemical factors used to amplify the HHC population affect the resulting iMNs, we compared the molecular and functional properties of iMNs generated in the 6F, DD, and DDRR conditions. First, we evaluated the transcriptional state of Hb9::GFP+ cells from all three conditions by RNAseq to determine how the populations vary globally. Hb9::GFP+ cells collected from all three conditions clustered together compared to the starting cell population of MEFs (Fig. 5D). Cells from 6F and DDRR conditions were relatively similar, although we observed small variations between the two populations (Fig. 5D-F). The addition of DDRR led to the downregulation of twice as many genes as those that were upregulated (Fig. 5E). Consistent with the increased efficiency of reaching the terminal iMN state in DDRR condition, Hb9::GFP+ cells showed downregulation of genes associated with fibroblasts such as extracellular matrix organization and collagen metabolic processes compared to 6F only (Fig. 5F). DDRR upregulated regulatory targets involved in kinase activity, cell cycle, and migration that reflect processes expected to be modulated by overexpression of hRasV12. Taken together these data indicate that DDRR accelerates the transcriptional shift away from the fibroblast state without loss of fidelity to the induced motor neuron profile generated by 6F conditions.

Accelerating maturation of lineage-converted cells remains one of the preeminent challenges limiting translational studies (23). To determine if expanding the HHC population accelerates maturation of the resulting cells, we examined the morphological and electrophysiological properties of iMNs. In vivo, neurons adopt different morphologies with varying polarity (e.g. unipolar, bipolar, multipolar). These morphologies are unique to their function and developmental window and impacts signal processing (35,36). Mature spinal motor neurons are multipolar (35-37). We found that inclusion of DD significantly increased the percentage of multipolar neurons generated for both mouse and human derived iMNs (Fig. 5G, H). These results suggest that inclusion of DD increases or accelerates the maturation of motor neurons generated by direct lineage conversion.

The chief function of mature motor neurons involves receiving and transmitting electrophysiological signals. Given that neuronal morphology influences electrophysiological behaviors (36,37), we next examined how DD impacted electrophysiological function of human iMNs. Using patch clamp electrophysiology, we measured sodium and potassium currents in iMNs generated with and without p53DD. DD iMNs displayed significantly larger sodium and potassium currents compared to 6F iMNs (Fig. S11A, B). Similarly, DD iMNs displayed faster, more mature action potentials (AP) (Fig S11C, D). Both metrics indicate more mature ion channel organization across the surface of DD iMNs. Finally, we examined the ability of mouse iMNs to adapt to repetitive stimulation (35). Upon repetitive stimulation, mature neurons display spike-frequency adaptation (SFA), an increasing time interval between spikes during an evoked series of action potentials, quantified as a ratio of the time interval between the last two APs of a series to that of the first two APs (35). Unlike 6F iMNs, DD iMNs displayed more robust spike frequency adaptation, with an SFA ratio several-fold higher than that of 6F iMNs (Fig. 5I-K). The SFA ratio from DD iMNs measured 5x higher than that reported for iPSC-derived iMNs with prolonged culture maturation (35). Taken together, the morphological and electrophysiological data indicate that expanding HHCs during reprogramming results in the production of iMNs that possess greater functional maturity.

## Discussion

Although direct lineage conversion enables access to increasing numbers of cell types for translational studies, reprogramming remains a rare event. Our studies into the systems-level features of reprogramming have identified combined hypertranscription and hyperproliferation as a central driver of reprogramming. Our data showing that transcription factor-induced hypertranscription impairs proliferation suggests that an antagonistic relationship exists between transcription and proliferation. Consistent with this notion, we find that few cells possess the processing capacity to mediate high rates of both processes. Consequently, reprogramming remains a rare event, restricted to privileged cells with high transcriptional and proliferative capacity. We find that HHCs contribute to the majority of reprogramming events and reprogram at near-deterministic rates. By introducing chemical and genetic perturbations that expand capacity, we can increase the population of HHCs, boosting the reprogramming rate and extending conversion to otherwise unreprogrammable populations. Through single-cell RNA-seq, we identified topoisomerase expression as a key parameter modulating a cell’s ability to sustain combined hypertranscription and hyperproliferation and undergo reprogramming. Topoisomerases mediate collisions between transcriptional and DNA-replicative machinery as well as resolve supercoils introduced by both processes. Our data suggest a model in which limits in reprogramming arise from tradeoffs between antagonism between transcription and replication rates that results from topoisomerase-related mechanisms such as torsional strain (Fig. 6). When replication and transcription exceed the cell’s capacity to resolve topological tangles and DNA breaks, both replication and transcription stall, retarding reprogramming processes. In the absence of perturbations that enable hyperproliferation and sustain hypertranscription, few cells possess the cellular machinery to balance the dynamic demands of rapid proliferation and hypertranscription. Increasing expression of topoisomerases expands the cell’s ability to mediate conflict between these two processes, enabling robust cellular reprogramming. Our data suggest that Top1 principally promotes transcription and Top2a promotes replication. Supporting the competing, dynamic demands of transcription and replication requires balanced expression of each topoisomerase.

**Figure 6.**
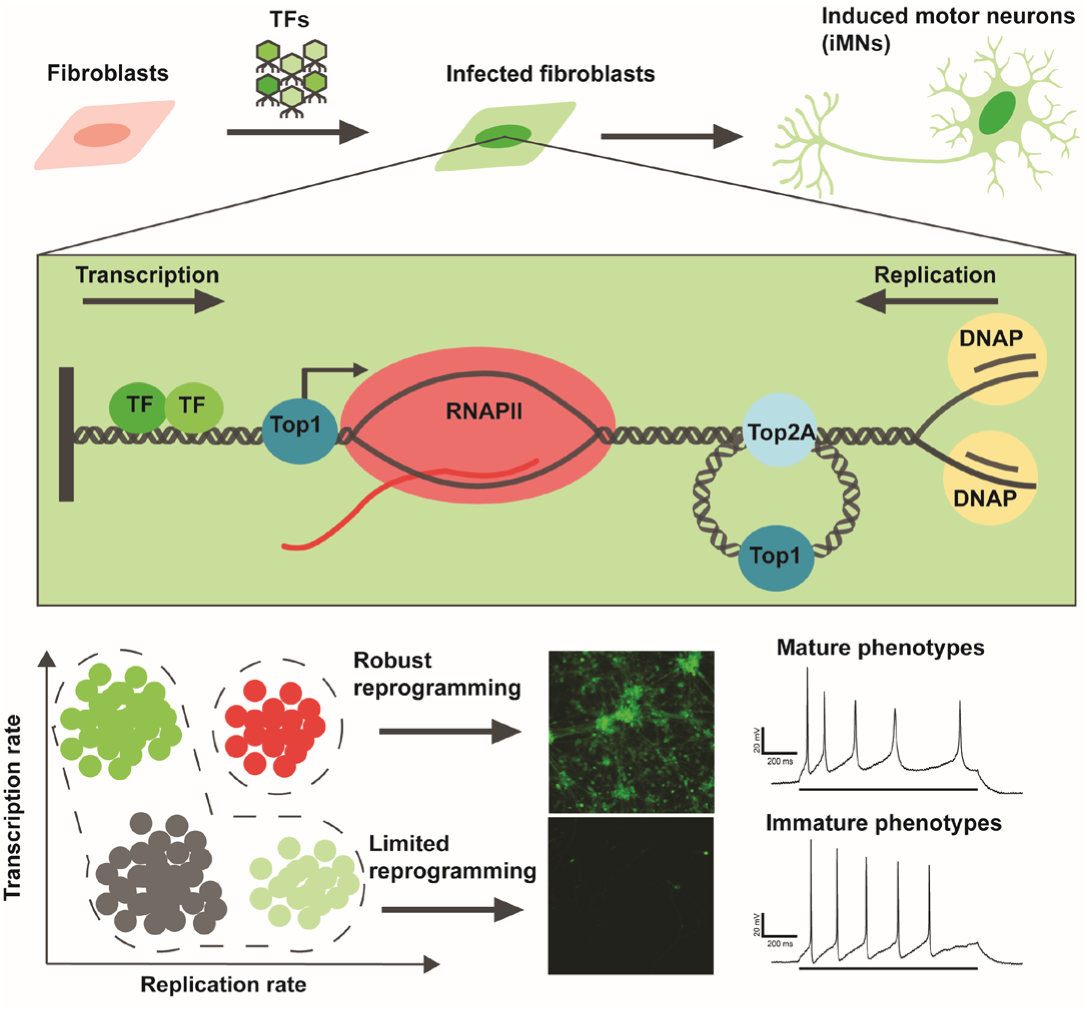
Model of topoisomerase-mediated reprogramming through hypertranscribing, hyperproliferating cells. Introduction of the reprogramming factors (TFs) to fibroblasts induces transcription and reduces cell cycle rate for most cells. For cells that continue to rapidly cycle, few sustain high rates of transcription. Increased expression of topoisomerases, Top1 and Top2a, which resolve supercoils and mediate conflicts in transcription and replication machinery, support the rare population of hypertranscribing, hyperproliferating (HHCs). HHCs and reprogram at near-deterministic rates to generate functionally mature cell type such as induced motor neurons (iMNs).

It is perhaps surprising that hyperproliferation stimulates reprogramming into post-mitotic lineages. However, previous work in E. coli indicates that rapid cell cycle promotes state switching (38). Therefore, rapid proliferation may be a general motif to facilitate cell fate transitions, while slower proliferation favors stable maintenance of cellular identity. Proliferation may facilitate the transcriptional shift away from the fibroblast identity. For example, macrophages have been shown to stabilize commitment to the myeloid lineage by lengthening their cell cycle as they exit the progenitor phase, leading to the accumulation of highly stable PU.1 (39). Differences in cell cycle rate most significantly impact the concentrations of highly stable molecular species. In the context of conversion, rapid replication may facilitate dilution of highly stable mRNAs and proteins (e.g. collagens) that may limit full adoption of an alternative identity.

Attaining a mature somatic cell state remains a major difficulty limiting the translational utility of reprogrammed or stem cell-derived cells (8,23). We show that cells that exhibit combined hypertranscription and hyperproliferation are capable of achieving greater functional maturity in the reprogrammed state. We hypothesize that enhanced maturity results from a greater ability of converting cells to remodel their GRN and proteome due to better dilution of the starting cell components, allowing more complete transition to the target state. Our study suggests that the biophysical properties of cells provide a formidable barrier to cellular transitions and inhibit maturation of in vitro derived cell types. We demonstrate that by increasing the cell’s capacity to balance tradeoffs during conversion we surmount maturity barriers.

While we have considered the synthetic transition of fibroblasts to motor neurons and other post-mitotic cells, our findings raise questions about transition of healthy cells to pathological states such as cancer. The genetic mutants we have employed to promote reprogramming are known to contribute to oncogenesis. The mechanisms that we have uncovered, such as replication prior to differentiation and hypertranscription, are recognized motifs in development (40,41). The overlap in developmental and oncogenic processes suggest central mechanisms for promoting transitions of cellular identity. Observing these stereotypical patterns of transitions in the synthetic context of reprogramming strengthens the hypothesis that healthy cells co-opt developmental processes in transit to their pathological state. Our data suggest that topological stress is a primary barrier to cellular reprogramming. In the context of cancer, our data would suggest that topological tangles act as genomic stabilizers of cellular identity, buffering cells against pathological transitions. Cancerous cells commonly express high levels of topoisomerases and topoisomerase inhibitors represent some of the front-line chemotherapy agents. The model of cellular conversion may provide a useful system to screen molecules that effectively block pathological transitions while preserving cells that maintain non-pathogenic states. Small molecules and cocktails that block the development of the HHC state may illuminate new therapeutic agents for targeting treatment to highly pathogenic cell states.

Our observations highlight a challenge for synthetic circuits integrated into large transcriptional networks. While significant efforts have been devoted to the logical design of enhanced synthetic circuitry, less is understood regarding how cellular hardware and the three-dimensional structure of genetic elements may impose fundamental limitations on integrated circuits. Our work suggests that the enhanced design of reprogramming vectors to account for limitations in cellular hardware may improve the predictability and determinism of reprogramming. Vectors designed to scale with the capacity of individual cells may more reliably enable reprogramming by limiting unnecessary transcriptional strain on the genome. Simple selection of promoters to regulate transcription-factor expression may be sufficient to improve expression-scaling from transgenic constructs. In addition, encoding factors that suppress the starting cell gene regulatory network may reduce the levels of transcription needed for successful reprogramming and therefore increase efficiency. Finally, developing principles that define how the three-dimensional context of the genome impacts integrated gene circuits will expand our ability to generate predictable cellular behaviors and improve cellular engineering. By expanding our understanding of how cells structure and integrate information across multiple molecular systems, molecular systems biology will continue to improve how we build cellular models and develop therapeutic strategies.

## Acknowledgments

We thank the NINDS Biorepository at Coriell Institute for providing the following cell lines for this study: ND39023. We thank the lab members of the Ichida lab for their useful feedback, and the University of Southern California Department of Animal Resources staff for their outstanding care.

## Funding

KNB was supported in part by CIRM Predoctoral Training Grant TG2-01161. KEG was supported by a Kirschstein-NRSA Postdoctoral Fellowship 5F32NS092417-03. This work was made possible by NIH grants R00NS077435, R01NS097850, and 5R01DC015530, US Department of Defense grant W81XWH-15-1-0187, and grants from the Donald E. and Delia B. Baxter Foundation, the Alzheimer’s Drug Discovery Foundation and the Association for Frontotemporal Degeneration, the Harrington Discovery Institute, the Tau Consortium, the Pape Adams Foundation, the Frick Foundation for ALS Research, the Muscular Dystrophy Association, the New York Stem Cell Foundation, the USC Keck School of Medicine Regenerative Medicine Initiative, the USC Broad Innovation Award, and the Southern California Clinical and Translational Science Institute to JKI. JKI is a New York Stem Cell Foundation-Robertson Investigator and a Richard N. Merkin Scholar. KK and BZ are supported by the NS090904, the NS100459, and the Foundation Leducq Transatlantic Network of Excellence for the Study of Perivascular Spaces in Small Vessel Disease reference no. 16 CVD 05.

## Author contributions

KNB and KEG designed and planned the experiments, performed the experiments, supervised experiments, analyzed the data, and wrote the manuscript. KK designed, performed, and analyzed all electrophysiology experiments and BZ supervised the work. YL designed and performed single-cell tracking experiments. MZ and BQ collected single-cells for qRT-PCR experiments and RC supervised the work. JKI designed and supervised experiments, and wrote the manuscript. All authors agreed with the final version of the manuscript.

## Competing interests

JKI is a co-founder of AcuraStem. JKI declares that he is bound by confidentiality agreements that prevent him from disclosing details of his financial interests in this work.

## Data and materials availability

The datasets used and/or analyzed during the current study are available from the corresponding author on reasonable request.

## Supplementary Materials

Materials and Methods

Figures S1-S11

Tables S1

External Databases S1-S4

References (42-48)

